# Identifying Isl1 genetic lineage in the developing olfactory system and in GnRH-1 neurons

**DOI:** 10.1101/2020.08.31.276360

**Authors:** Ed Zandro M. Taroc, Raghu Ram Katreddi, Paolo E. Forni

## Abstract

During embryonic development, symmetric ectodermal thickenings (olfactory placodes) give rise to several cell types that comprise the olfactory system, such as those that form the terminal nerve ganglion (TN), gonadotropin releasing hormone-1 (GnRH-1) neurons and other migratory neurons in rodents. Even though the genetic heterogeneity among these cell types are documented, unidentified cell populations arising from the olfactory placode remain. One candidate to identify placodal derived neurons in the developing nasal area is the transcription factor Isl1, which was recently identified in GnRH-3 neurons of the terminal nerve in fish, as well as expression in neurons of the nasal migratory mass. Here, we analyzed the Isl1 genetic lineage in chemosensory neuronal populations in the nasal area and migratory GnRH-1 neurons in mice using *in-situ* hybridization, immunolabeling a Tamoxifen inducible Isl1Cre^ERT^ and a constitutive Isl1^Cre^ knock-in mouse lines. In addition, we also performed conditional Isl1 ablation in developing GnRH neurons. We found Isl1 lineage across non sensory cells of the respiratory epithelium and sustentacular cells of OE and VNO. We identified a population of transient embryonic Isl1+ neurons in the olfactory epithelium and sparse Isl1+ neurons in postnatal VNO. Isl1 is expressed in almost all GnRH neurons and in approximately half of the other neuron populations in the Migratory Mass. However, Isl1 conditional ablation alone does not significantly compromise GnRH-1 neuronal migration or GnRH-1 expression, suggesting compensatory mechanisms. Further studies will elucidate the functional and mechanistic role of Isl1 in development of migratory endocrine neurons.

## 1) Introduction

Cranial placodes are specialized regions of ectoderm, that give rise to the pituitary gland, sensory organs and ganglia of the vertebrate head [1; 2]. They form as a result of specific expression patterns of transcription factors in pre-placodal ectoderm surrounding the anterior neural plate [2; 3]. In mice, the olfactory placodes (OPs) are morphologically identifiable at embryonic day 9.5 (E9.5), as a bilateral ectoderm thickening in the antero-lateral region of the head (fig1). The transcription factors Oct1, Sox2, Pou2f1, Pax6, Eya1, Six1, Six4, Ngnr1,Hes1, Hes5, MASH1/Ascl1, Ngn1 and Gli3 have been identified to control various steps of olfactory placode induction/formation, neurogenesis, neuronal development, and expression of olfactory specific genes [4; 5; 6; 7; 8; 9; 10].

**Fig. 1).**
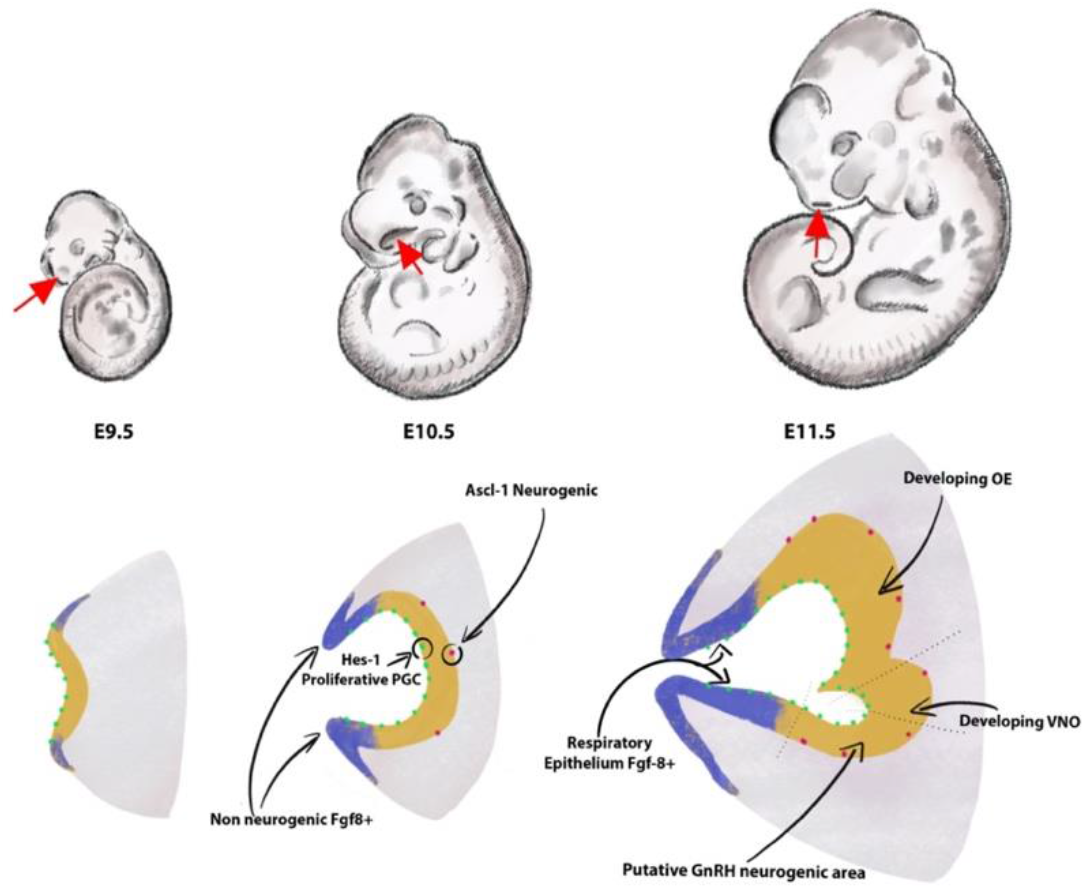
Formation of the olfactory pit. Drawings depict developing embryos between E9.5 and E11.5. The frontonasal areas where the olfactory placodes/pits form are indicated with red arrows. Cartoons in the lower row illustrate the olfactory placode ingression between E9.5 and E10.5 (chronal view) and a parasagittal view of the olfactory pit at E11.5. The lateral and rostral portions of the invaginating olfactory placode (blue) act as the main source of FGF8. These Fgf8+ areas progressively differentiate into respiratory epithelium. In the neurogenesis portion of the olfactory placode (yellow), proliferative Hes-1+ progenitors (green) give rise over time to neurogenic Ascl-1 neuronal progenitors. Arrows show the vomeronasal organ (OE) and putative neurogenic area of the GnRHns at E11.5 in the developing olfactory epithelium (OE),. The size of the pit and cells is arbitrary and does not reflect natural proportions

The olfactory pit of vertebrates generates neurogenic and non-neurogenic progenitors. The non-neurogenic portions of the olfactory pit give rise to the respiratory epithelium, which is located in the rostral olfactory pit, while the neurogenic portions give rise to specific neuronal and glial/support cells (Fig.1,2). The respiratory epithelium, which is the main source of Fgf8, controls the development of the neural crest derived nasal mesenchyme and bones [11]. Neurogenic waves in the olfactory pit of mice first gives rise to mostly migratory neurons [11; 12] such as early pioneer olfactory neurons, neurons of the terminal nerve, including Gonadotropin releasing hormone neurons (GnRHns), NPY positive migratory neurons, and other neurons with unknown identity and function or neurons of the migratory mass (MM) [13; 14; 15; 16]. The nasal area of mice contains several other neuronal cell types that include specialized olfactory sensory neurons such as the guanylyl cyclase-D (GC-D) neurons of the necklace olfactory system [17; 18; 19], microvillar cells (MVCs)[20], sensory neurons of the septal organ (SO) [21], the Grueneberg ganglion (GG) [22; 23; 24; 25; 26], and cells forming the terminal nerve ganglion (TN) [27; 28; 29; 30; 31; 32; 33; 34] including the GnRH-1 ns [14; 15]. The mechanisms and molecules that drive progenitors of the developing olfactory pit into early migratory cells types remains largely unknown.

**Fig. 2).**
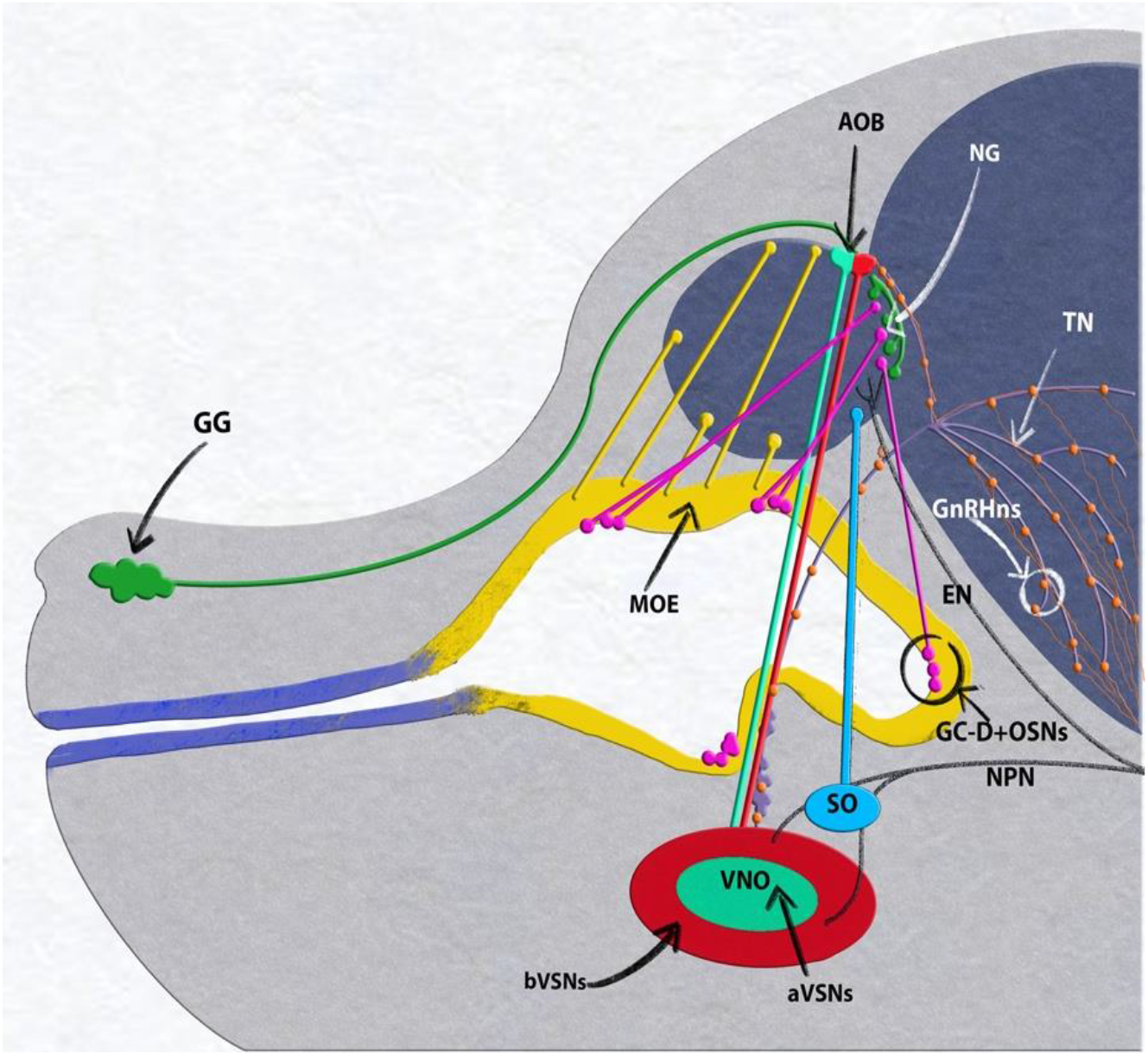
E15.5 mouse embryo, neuronal populations in the nasal area. Grueneberg ganglion (GG, green) and GC-D+/necklace (OSNs, magenta) project to the necklace glomeruli (NG). Olfactory neurons of the main olfactory epithelium (MOE, yellow) projecting to the main olfactory bulb (MOB). Vomeronasal organ (VNO), composed of basal (bVSNs, green) and apical (a-VSNs, red), projecting to anterior and posterior portions of the accessory olfactory bulb (AOB). Septal Organ (SO, light blue) projecting to the ventral OB. Terminal nerve (TN, purple) projecting to the basal forebrain, GnRHns, orange, scattered along the terminal nerve fibers from the vomeronasal area to the basal forebrain. Trigeminal nasopalatine (NPN) and ethmoidal nerve projections (EN) drawn in dark gray.

Between E 9.5 and E10.5, the OPs begin to ingress to form the olfactory pits that give rise to various cellular components of the nose during development [6; 35]. The olfactory pit is largely populated by proliferative progenitors expressing the transcription factors Hes1+/Hes5. These early progenitors undergo mitosis in the apical portion of the epithelium and give rise over time to neurogenic progenitors positive for the bHLH transcription factor Ascl-1 [35; 36; 37; 38]. Gli3 influences proliferative Hes1+ positive progenitors to give rise to Ascl-1+ neurogenic progenitors that eventually generate olfactory sensory neurons (OSNs) and vomeronasal sensory neurons (VSNs) [36; 37]. Notably, Gli3 loss of function does not prevent neurogenesis of GnRH-1 neurons (GnRH-1ns), suggesting that these have a distinct lineage from the olfactory neurons.

GnRH-1ns are the initial neuronal population to form in the developing olfactory pit [12; 14]. During embryonic development, GnRH1ns migrate along the axons of the terminal nerve [28] from the nasal area to the basal forebrain. Once in brain the GnRH-1ns control the release of gonadotropins from the pituitary gland [39]. Aberrant development or function of GnRH-1ns causes hypogonadotropic hypogonadism (HH) [40], that can lead to a spectrum of reproductive disorders [41]. GnRH-1 neuronal migration strictly depends on correct development and maturation of the neural crest derived olfactory ensheathing cells [42; 43; 44; 45]. Cre genetic lineage tracing and bac transgenic reporters have revealed extensive genetic heterogeneity among neurons that originate in the olfactory area of mice including the GnRHns, suggesting a potential integration of neural crest cells in the developing OP [28; 46; 47; 48; 49]. However, the mechanisms that give rise to the GnRH neurons and the required transcription factors for these regulatory processes remain unresolved.

The LIM-homeodomain transcription factor Isl1 has been recently identified in subsets of neuronal derivatives forming from the olfactory placode [49; 50; 51]. Isl1/2 expression occurs in GnRH expressing neurons and other putative neurons of the terminal nerve [49; 50; 51; 52] in chick, zebrafish, mouse, and humans. In zebrafish [50], Isl1/2 is expressed in terminal nerve GnRH-3 neurons associated with the olfactory epithelium. These neuro-modulatory cells, which are non-migratory in fish, exert the same physiological function as migratory GnRH-1 neurons in mice [39]. The terminal nerve and migratory mass in birds contains heterogeneous populations of cells that 1) express GnRH-1 but are negative for Isl1/2 and Lhx2, 2) cells only expressing Lhx2, 3) cells only positive for Isl1, and 4) cells that co-express Lhx2, Isl1/2, GnRH-1 [51]. In birds, cells from the terminal nerve ganglion (TN) are also positive for Isl1 and Lhx2 that differ from the GnRH-1ns. A third study in mouse [49] also confirmed Isl1/2 immuno-reactivity GnRH-1ns at early stages of GnRH neuronal development (E11.5). A subset of GnRH-1ns, positive for Wnt1Cre genetic lineage [46], were negative for Isl1 immunoreactivity.

One prediction is that Wnt1Cre tracing, a controversial neural crest marker, and Isl1 immunoreactivity define two distinct embryonic lineages for GnRH-1ns. The rival hypothesis is that the Wnt1Cre positive subpopulation of GnRH-1ns differ in its timing and/or expression levels of Isl1 from the majority of GnRH-1ns [46; 49]. Understanding the molecular cascade that establishes the genetic lineage of different cell types is a fundamental step to identify potential cellular differences between cell types and to unravel the molecular mechanisms underlying neurodevelopmental pathologies. Here, we examined which regions in the OP express Isl1/2 and and analyzed Isl1 genetic lineage in different neuronal types in the nasal area of mice. We further generated Isl1 conditional KO mice to study the role of Isl1 in GnRH development.

## MATERIAL AND METHODS

### Animals

Isl1^flox^(Isl1tm2Sev/J) [53]; Isl1^CreERT^(Isl1tm1(cre/Esr1*)Krc/SevJ) [54], Isl1Cre (Isl1tm1(cre)Sev/J) [55] Rosa26tdTomato (B6.Cg-Gt(ROSA)26Sortm14(CAG-tdTomato)Hze/J) [56] and GnRH-Cre (Gnrh1-cre 1Dlc/J) [57] mouse lines were all purchased from JAX. Mice of either sex were used for ISH and IHC experiments. All experiments involving mice were approved by the University at Albany Institutional Animal Care and Use Committee (IACUC).

### Tamoxifen treatment

Tamoxifen (Sigma-Aldrich), CAS # 10540-29-1 was dissolved in Corn Oil. Tamoxifen was administered once via intraperitoneal injection at indicated developmental stages. Tamoxifen was administered at 180mg/Kg based on the weight of the pregnant dam on the day of injection.

### Tissue preparation

Embryos were collected from time-mated dams where the emergence of the copulation plug was taken as E0.5. Collected embryos were immersion-fixed in 3.7% Formaldehyde/PBS at 4°C for 3-5 hours. Postnatal animals were perfused with 3.7% Formaldehyde/PBS. All samples were then cryoprotected in 30% sucrose overnight, then frozen in O.C.T (Tissue-TeK) and stored at −80°C. Samples were cryosectioned using CM3050S Leica cryostat and collected on Superfrost plus slides (VWR) at 14μm thickness for embryo’s and 16μm for post-natal tissue.

### Immunohistochemistry

Primary antibodies and dilutions used in this study were: rabbit-α-Isl1(1:1000, Abcam), mouse-α-Isl1/2 (1:100, DSHB), SW rabbit-α-GnRH-1 (1:6000, Susan Wray, NIH), goat-α olfactory marker protein (1:4000, WAKO), rabbit-α-phospho-Histone-H3 (1:400, Cell Signaling), goat-α-Sox2 (1:400, R & D systems), rat-α-phospho-Histone-H3 (1:500, Abcam), mouse-α-Ki67 (1:500, Cell Signaling), rabbit-α-DsRed (1:1000, Clontech), goat-DsRed (1:1000, Rockland), HuC/D 8ug/ml (Molecular Probes). Antigen retrieval was performed in a citrate buffer prior to incubation with rabbit-α-Isl1, mouse-α-Isl1/2, rabbit-α-phospho-Histone-H3, rat-α-phospho-Histone-H3, mouse-α-Ki67. For immunoperoxidase staining procedures, slides were processed using standard protocols [11] and staining was visualized (Vectastain ABC Kit, Vector) using diaminobenzidine (DAB) in a glucose solution containing glucose oxidase to generate hydrogen peroxide; sections were counterstained with methyl green. For immunofluorescence, species-appropriate secondary antibodies were conjugated with Alexa-488, Alexa-594, or Alexa-568 (Molecular Probes and Jackson Laboratories) as specified in the legends. Sections were counterstained with 4′,6′-diamidino-2-phenylindole (1:3000; Sigma-Aldrich) and coverslips were mounted with Fluoro Gel (Electron Microscopy Services). Confocal microscopy pictures were taken on a Zeiss LSM 710 microscope. Epifluorescence pictures were taken on a Leica DM4000 B LED fluorescence microscope equipped with a Leica DFC310 FX camera. Images were further analyzed using FIJ/ImageJ software. Each staining was replicated on at least three different animals for each genotype.

### In situ hybridization

Digoxigenin-labeled cRNA probes were prepared by in vitro transcription (DIG RNA labeling kit; Roche Diagnostics) from the following templates: Isl1 from Allen Brain Atlas (Probe RP_080807_04_G12). In situ hybridization was performed as described [58] and visualized by immunostaining with an alkaline phosphatase conjugated anti-DIG (1:500), and NBT/BCIP developer solution (Roche Diagnostics).

### Cell quantifications

GnRH-1/Isl1 double IHC quantifications were performed at E11.5 and E15.5 parasagittal non-serial sections on a single slide using the Rabbit-α-GnRH-1 primary and Mouse-α-Isl1/2 primary antibodies. Embryonic co-localized cell counts on Isl1Cre or Isl1Cre^ERT^ lineage traced animals were done on E14.5 and E15.5 respectively parasagittal non-serial whole head sections within the nasal region on a single slide where we see co-localization of both markers indicated (DsRed/tdTomato GnRH-1, or HuCD). Post-natal OMP cell counts were done on P8 coronal nose sections. To get the density of mature OSNs positive for constitutive Isl1 lineage tracing at P8, number of OMP positive neurons colocalized for tdTom were counted in the caudal olfactory epithelium and calculated the area of the OMP positive cells within the traced epithelium. In VNO, to get the percentage of mature VSNs positive for constitutive Isl1 lineage tracing, firstly OMP positive cells colocalized for tdTom were counted. To get the total OMP positive cells in VNO section, density of OMP cells in smaller areas of VNO was determined and extended to the total area of OMP positive cells in the VNO.

GnRH-1 cell counts were performed on GnRH-1Cre/Islet1^flox/flox^, Isl1^CreERT/flox^, and control at E15.5 on two immunostained non-serial slides.

### Validation of conditional Isl1 knockout

The Isl1^flox^ animals have the lox-p sites flanking exon 4 of the Isl1 gene which codes for the homeodomain and consists of the amino acids in positions 181-240. To detect conditional loss of Isl1^flox^ in GnRHCre/Isl1^flox/flox^ mutants we performed immunolabeling using the mouse anti-Isl1/2 (DSHB) primary antibody. This antibody recognizes the amino acids 178-349 of the Isl1 protein and was previously shown to be unable to detect Isl1^flox^, (Isl1tm2Sev/J) mice, after Cre mediated recombination [53]. However, in Isl1^CreERT^ (Isl1tm1(cre/Esr1*)Krc/SevJ) which is null mouse model, the amino acids in positions 181-240 are maintained, therefore this null protein is still detectable in Isl1 null cells. Since Isl1^CreERT^ [54] and Isl1^flox^ alleles [53] have been targeted in different gene regions we could not validate recombination efficency of Isl1^CreERT^/^flox^ cKOs.

### Statistical Analyses

All statistical analyses were carried out using GraphPad Prism7 software. All cell counts were averaged ± standard error (SE) among animals of the same age and genotype. Means ± SEs were calculated on at least three animals per genotype. The statistical difference between genotypes and groups were determined using an unpaired student’s t-test.

## RESULTS

### Isl1 expression in proliferative progenitors and neurons during early development of the olfactory pit

Using in-situ hybridization at E11.5, we found Isl1 mRNA expression in the more caudal region of the putative developing olfactory epithelium and proximal to the developing VNO (Fig. 3 A). We previously described the latter as the region where GnRH-1ns likely form and become immunoreactive (Fig 3 B) [11]. By pairing immunohistochemistry against Isl1/2 and the mitotic marker phospho-histone-3 (PHH3) at E11.5, we found (Fig. 3 C,C’) some proliferative progenitors in the apical regions of the developing pit that were also positive for Isl1. Isl1/2 protein expression was consistent with the mRNA expression pattern (Fig 1A, C). However, strong Isl1/2 immunoreactivity appeared localized in sparse nuclei ventral to the developing VNO and the putative respiratory epithelium (Fig 1B-D). Immunolabeling with anti Isl1/2 and GnRH-1 at E11.5, E13.5 and E15.5 (Fig 3D-F’’’) indicated a dynamic Isl1/2 expression across GnRH-1ns [49]. In fact, we could identify at E11.5 approximately 89% (SE +/−3.33%) of GnRH-1+ cells positive for Isl1/2 immunoreactivity (Fig 3D-D”). However, GnRH-1ns were negative for Isl1/2 immunoreactivity in both nasal area and brain by E15.5.

**Figure 3.**
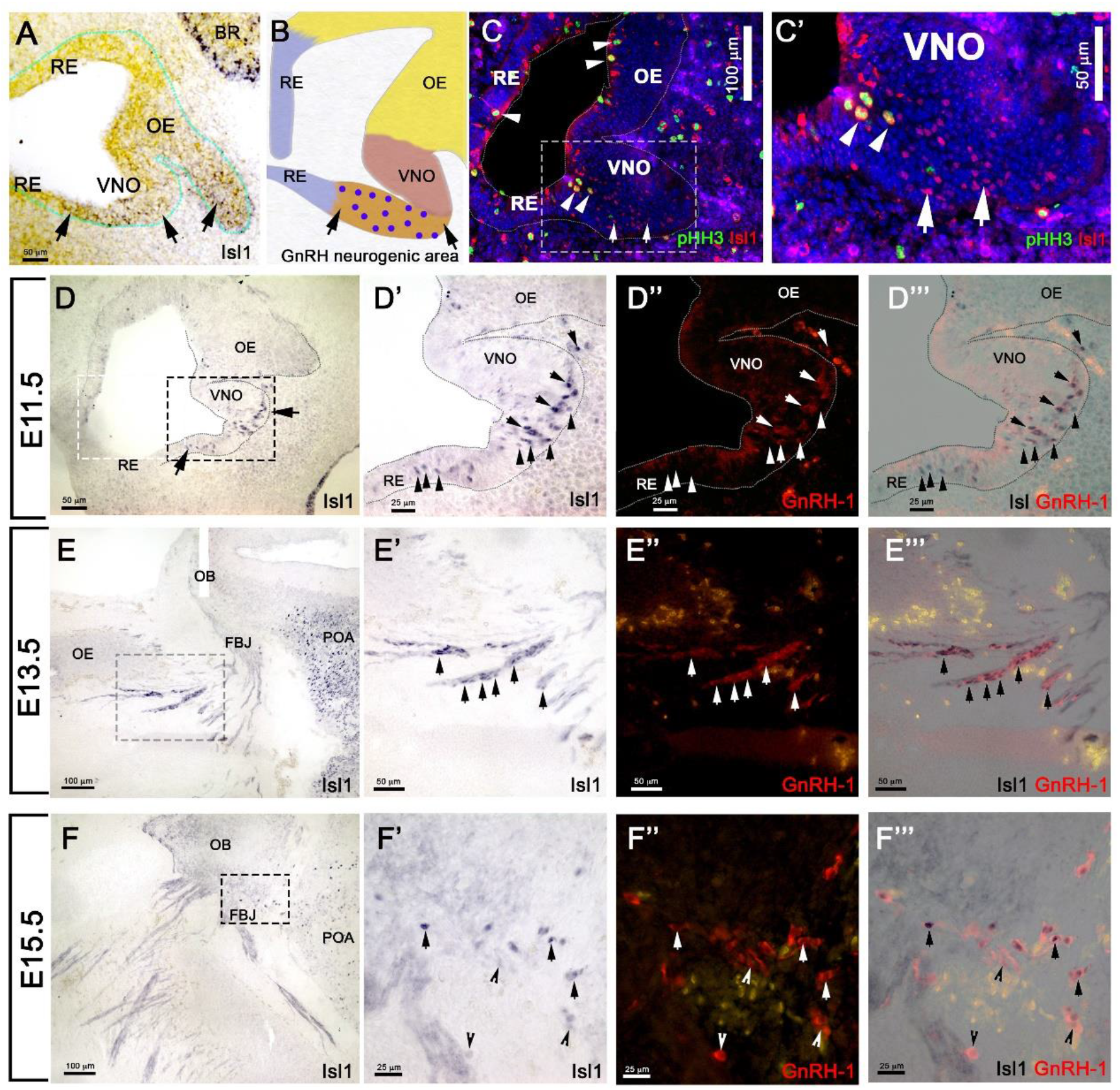
Isl1 expression in the OP is restricted to GnRH neurons and a few other cells types. A) E11.5 *In-situ* hybridization anti-Isl1 mRNA shows Isl1 expression in the ventral portion of the developing olfactory pit proximal to the VNO. Arrows indicate putative GnRH neurogenic area. Only spare cells positive for Ilset1 could be detected in the OE (arrowhead). B) Cartoon illustrating the putative areas giving rise to RE, OE, VNO and the putative GnRH neurogenic area. Blue dots indicate GnRHns. C,C’) Immunostaining anti-Isl1 (red) and the mitotic marker pHH3 (green) shows Ilset1 expression in sparse proliferative cells in the apical portion of the OP (arrowheads) and in cells in the putative GnRH neurogenic area ventral to the VNO (boxed) compare to A,B. D) Isl1 immunoreactivity in the putative GnRH neurogenic area (arrows, compare to A,B,C). C’-C”’) Isl1 expression (black) in newly formed GnRH neurons(arrows) and in GnRH-1 negative cells (arrowheads). E-F”’ Isl1 expression (black) in migrating GnRHns in the (arrows) in the nasal area at E13.5 (E-E”’) and in the brain at E15.5. (F-F’”). Olfactory epithelium (OE), Olfactory bulb (OB), Forebrain junction (FBJ), Preoptic area (POA).

### Differential lineage among olfactory, vomeronasal and GnRH-1 neurons

As we observed sparse and a heterogenous distribution of Isl1 mRNA and protein expression in the developing olfactory pit at E11.5 (Fig. 3) and found Isl1/2 protein expression with the proliferative cell marker PHH3, we decided to investigate which cells formed from the Isl1/2+ proliferative cells. We utilized a tamoxifen inducible Isl1Cre^ERT^ knock-in mouse line [54] mated with a sensitive Rosa-reporter [56]. Isl1Cre^ERT^ allows temporal control and restricts Cre mediated recombination to the time of Tamoxifen injection. Pregnant dams were treated with a single injection of Tamoxifen (180mg/Kg) at E11.5 and embryos were analyzed 4 days later (Fig. 4A). At E15.5, we observed Isl1 recombination in trigeminal nerve, inner ear, Rathke’s pouch and oral mucosa (Fig.4B,C) [59; 60; 61; 62]. We also found Isl1 tracing in the developing nasal area (Fig. 4D-E”) in sparse OMP+ sensory neurons of both the main olfactory and vomeronasal epithelium. Notably, Isl1Cre+ neurons positive for the pan neuronal marker HuC/D were found at the border between the respiratory epithelium and sensory epithelium (Fig. 4E-E’). We found extensive Cre recombination penetrance throughout the nasal respiratory epithelium (Fig. 4,D,D’).

**Fig. 4.**
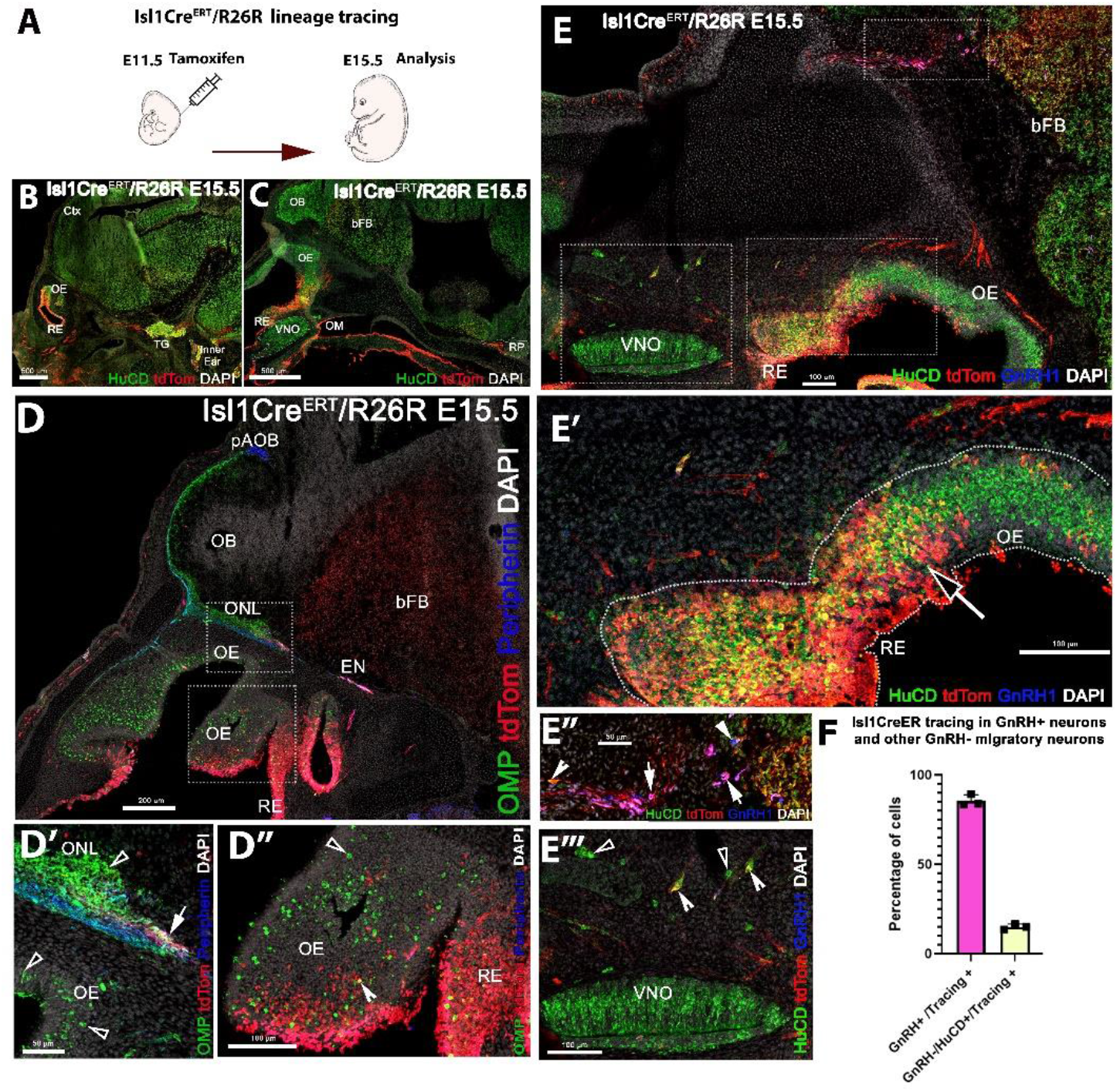
Temporally controlled Isl1 lineage tracing at E11.5 primarily highlights cells of the respiratory epithelium and GnRHns. A) Cartoon illustrating experiential strategy for Ilset-1CreERT/R26RTdTomato lineage tracing. Tamoxifen induction of Cre recombiantion at E11.5, analysis at E.15.5. B,C) HuC/D/DsRed double immunostaining shows recombination (tdTomato) in the Trigeminal Ganglion (TG), the inner ear as well as in cells of the Rathke’s Pouch (RP). D-D’’ In the developing nose recombination appears to be mostly restricted to the respiratory epithelium (RE) and Ethmoidal (EN) trigeminal projections. D-D” Anti tdTomato; OMP; Peripherin immunostaining showed that OMP positive olfactory projections of the main olfactory bulb. Peripherin positive projections to the accessory olfactory bulb appear negative for td Tomato expression. However, tdTomato positive fibers (D’ white arrow), likely belonging to the trigeminal/ethmoid nerve (See EN in D), appeared to project to the ventral OB. D”) OMP and tdTomato immunoreactivity showed nearly complete absence of OSNs positive for Isl1 tracing. Neurons in the olfactory area appeared mostly negative for Isl1 lineage D”, sparse OMP+ tdTomato + cells (Notched arrowhead) could be detected at the border between olfactory and respiratory epithelium, boxed area see D’. E-E’’ E’’’) Pan neuronal marker HuC/D shows neurons of the developing vomeronasal organ (VNO), mostly negative for Isl1 lineage (E’). HuC/D+ neurons at the border between OE and RE (E’) migratory neurons, GnRH-1ns (E”) and HuC/D+only (E”’) were positive for Isl1 tracing. F) Quantification of recombination in migratory GnRH-1+ and HuC/D GnRH-1 negative neurons.

We detected putative nasopalatine and ethmoidal trigeminal axons positive for Isl1 through-out the nasal area and tangential to the olfactory bulb (Fig 4B,C,D,D’). By analyzing the olfactory projections via olfactory marker protein (OMP) and Peripherin immunostaining in combination with anti tdTomato staining, we confirmed that Isl1Cre recombination occurred in neurons that appeared to bundle with the olfactory neurons. Notably, most Isl1Cre^ERT^ traced axons did not appear to project to the main or accessory olfactory bulb (Fig. 4D,D’). Analysis of the olfactory epithelium showed a very sparse colocalization between OMP and Isl1Cre mediated recombination. However, immunostaining against the pan neuronal marker HuC/D showed the presence of Isl1+ neurons within the developing olfactory epithelia (Fig. E, E’). Triple immunostaining against GnRH-1, tdTomato and the pan neuronal protein HuC/D showed that temporally controlled Isl1Cre^ERT^ recombination at E11.5 yielded 85% (SE +/−1.88%) of GnRH-1ns were positive for tracing, while only 15%(SE +/−0.91) of the migratory (HuC/D+/GnRH-) cells were Isl1 positive. These data suggest that Isl1 expression at E11.5 (Fig. 3 A-C) is mostly limited to progenitors or precursors of GnRH-1ns and the subsets of other neurons of the migratory mass.

### Analysis of Constitutive Isl1 Cre genetic lineage tracing at E14.5

Tamoxifen controlled Cre recombination at E11.5 indicated that few neuronal progenitors and sparse neurons in the OE and VNO were positive for Isl1 as the olfactory pit forms, while the respiratory epithelium, oral mucosa and the majority of the GnRH-1ns resulted expressing Isl1. To further perform a temporally unbiassed lineage tracing for Isl1, we exploited a traditional Isl1Cre knock-in mouse line. In these mice recombination is expected to occur whenever Isl1 is expressed [55]. We performed a Cre/Rosa reporter lineage analysis Isl1^Cre+/−^/R26^tdTomato+/−^ at E14.5 and P8. After temporally controlled recombination (Fig.4), constitutively active Isl1 Cre at E14.5 (Fig.5) showed recombination in the trigeminal nerve, cells of the inner ear [63], the Rathke’s pouch forming the anterior pituitary gland [64], the oral mucosa and the nasal respiratory epithelium.

**Fig. 5.**
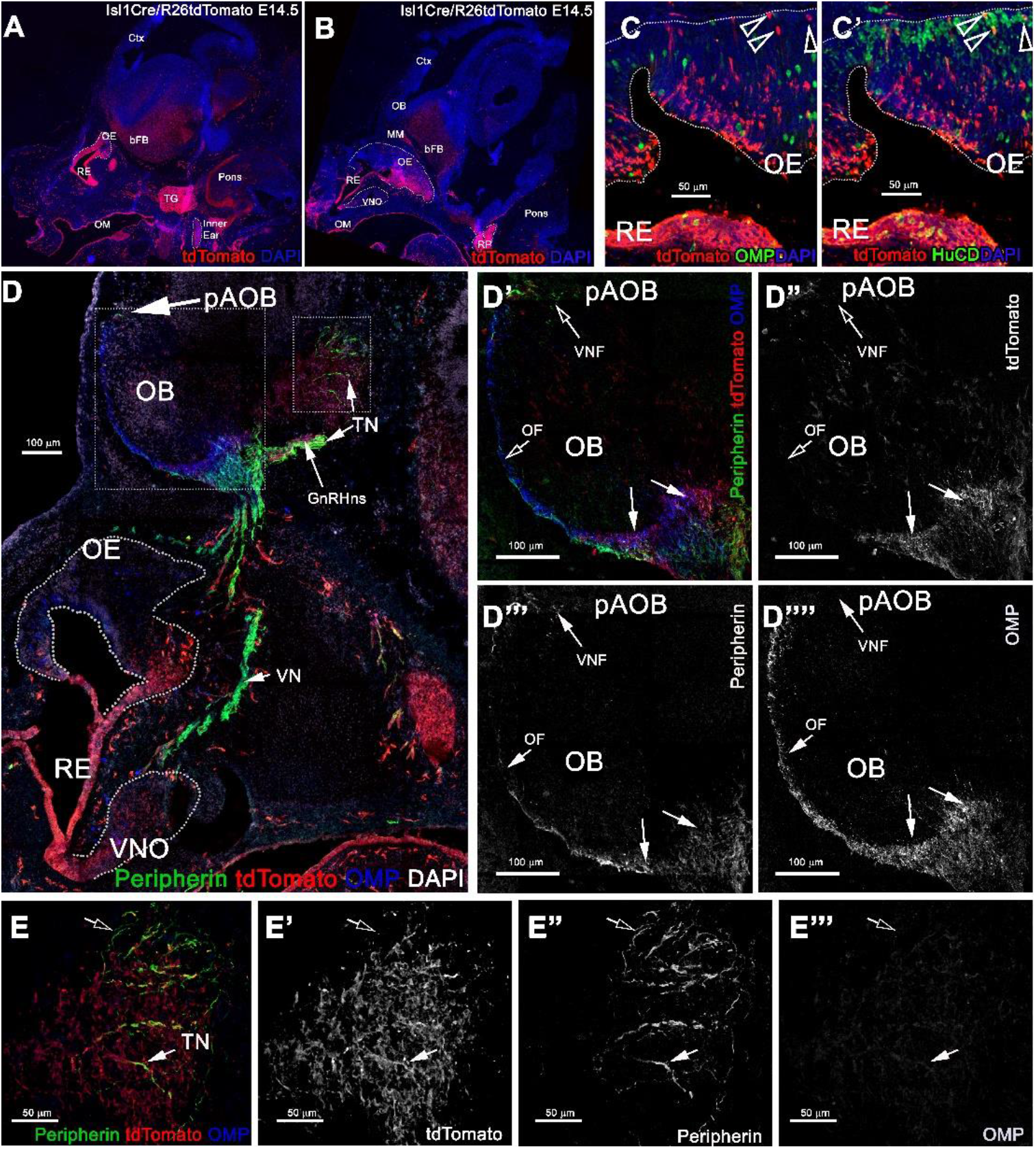
Constitutive Isl1 lineage tracing highlights: sparse neurons the OE, VNO, neuronal projections to the ventral OB, GnRH neurons and some TN fibers. A-B) Immunostaining against tdTomato of Isl1Cre/R26tdTomato lineage traced embryo’s at E14.5, showing recombination the trigeminal ganglion (TG), inner ear, Rathke’s pouch (RP), the oral mucosa (OM), Respiratory epithelium (RE) and in sparse cells in the OE and VNO. C) Anti-tdTomato; OMP and C’) anti td Tomato; HuC/D stainings shows that though almost all the the OMP+ cells in the were negative for tracing (arrowheads) (C), those were neurons positive for HuC/D (arrowhead). D-E’’’) Immunostaining against tdTomato, OMP, and Peripherin to highlight the projections of the developing olfactory system reveal that Isl1 tracing (D’-D’’’’) is mostly absent in the OMP+ olfactory fibers (OF) and Peripherin+ vomeronasal fibers (VNF) innervating the olfactory bulb (OB). GnRH-1 and other cells of the migratory mass positive for the tracing were detected invading the brain along the terminal nerve (TN). E-E”’ Peripherin positive TN fibers in the brain. Some fibers appear positive for the tracing (solid arrow) while other Peripherin fibers appear to be negative (empty arrow).

During gross visual observation, we noticed that more cells positive for Isl1 lineage occurred in both olfactory and vomeronasal epithelia compared to that after temporally controlled Cre activation (Compared Fig.5 and Fig.4).

### Isl1 Cre recombination in GnRH and terminal nerve neurons

At E14.5 we performed OMP, Peripherin and tdTomato immunostaining. We observed labeled olfactory and terminal axonal projections (Fig. 5 D-E’’’). In the olfactory system, we noticed that Isl1 Cre tracing could be seen in fibers forming bundles with the olfactory neurons after temporally controlled Isl1 Cre recombination (Fig. 5 D’-D’’’). However, most axons positive for recombination at this stage appeared to project only to the ventral portions of the olfactory bulb with some verging on invading the basal forebrain brain (Fig.5 E-E’’’’). The terminal nerve enters the brain ventral to the olfactory bulb [28]. Fibers of the putative terminal nerve associated with Isl1+ migratory GnRH-1ns. Using immunostaining against peripherin and tdTomato, we visualized Isl1+ positive cell somas and fibers of GnRH-1ns migrating on peripherin positive axons of the TN. Some putative TN’s axons showed sparse positivity for tracing (Fig.5 E-E’’’’). To confirm Isl1 Cre tracing in cells of the terminal nerve, we immunostained against the antigen Robo3, which is selectively expressed by these cells [28]. Robo3/tdTomato staining confirmed the presence of cell bodies from putative TN cells, proximal and within the developing vomeronasal organ, some of these appeared to be positive for Isl1 lineage. We also observed TN fibers, positive for Robo3, in the brain (data not shown). GnRH-1, tdTomato and HuC/D (Fig. 6 D-D’’’) immunostaining confirmed Isl1 lineage in virtually all GnRH-1ns (95%(SE +/−0.68), while 55% (SE +/−4.63) of the HuC/D migratory neurons that were negative for GnRH-1, were actually positive for Islet1 tracing (Fig. 6 E).

**Fig. 6.**
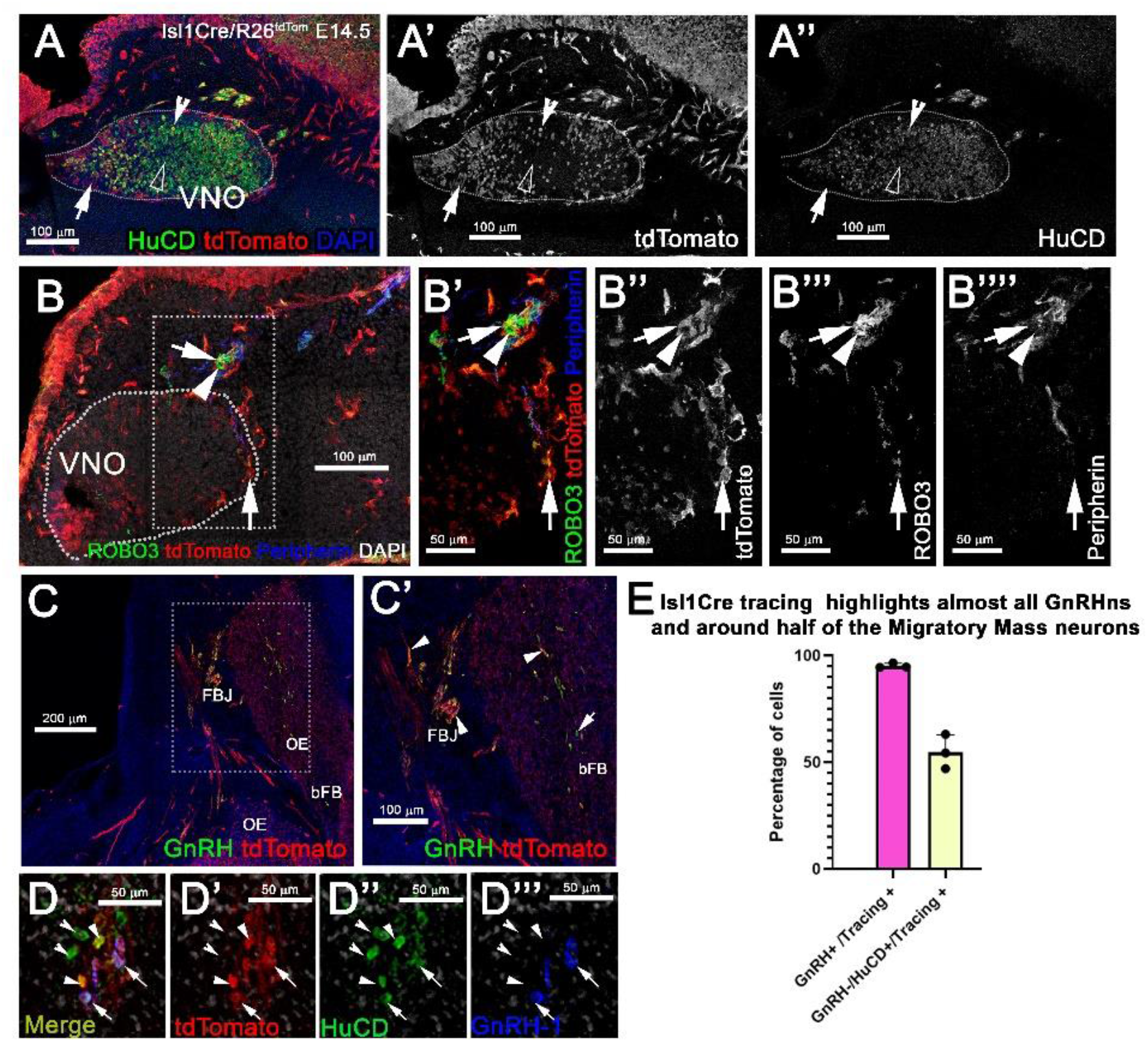
Isl1 constitutive lineage tracing highlights GnRHns, some cell bodies of the TN, and other migratory neurons. A-A’’) HuC/D and tdTomato immunostaining of Isl1Cre^+/−^/R26tdTomato VNO at E14.5 show some neurons within the VNO positive (empty arrowhead) and negative (notched arrowhead) for Isl1 recombination. Population of tdTomato+ non-neuronal cells positive for tracing (arrow). B-B’’’) Anti-Robo3; tdTomato and Peripherin labeling shows that a portion of putative terminal nerve cell bodies located within and proximal to the VNO are highlighted by Isl1 recombination (arrow) while a portion do not (arrowhead). C-C’) GnRH and tdTomato staining shows recombination in the majority of the GnRHns (arrowheads). D-D’’) shows recombination in ~ half of the (GnRH-;HuC/D+) migratory neurons (arrows. E) Quantification of Isl1Cre recombination in migratory GnRH+ and HuC/D GnRH-1 negative neurons

### Postnatal analyses of Isl1Cre lineage in the nasal cavity

During embryonic development, we observed Isl1 recombination in neurons in both olfactory and vomeronasal epithelia. So, we analyzed the pattern of constitutive Isl1 Cre recombination in postnatal animals. We performed double immunostaining against tdTomato and OMP at postnatal day 8 and noticed that most olfactory epithelium neurons did not express OMP and Isl1. However, we did find sparse neurons positive for Isl1 tracing in the most caudal ethmoid turbinates (0.12 (+/− 0.03 SEM) cells/1000μm^2^). Isl1Cre positive sustentacular cells were found close to the border between the respiratory epithelium and sensory olfactory epithelium in the ventral-lateral zones of the OE and along the nasal septum (S) in all the sections from rostral to caudal. At visual gross observation, we noticed more pronounced Isl1+ lineage in sustentacular cells in the caudal OE along the lateral ethmoid turbinates. These data imply spatial/regional heterogeneity in gene expression among sustentacular cells [65].

Analysis of postnatal vomeronasal epithelia showed a different scenario from that in the OE. Immunostaining against TdTom indicated Isl1 tracing in both non-sensory epithelium (NSE) and sensory epithelium of the VNO. In the sensory epithelium, staining against OMP and tdTom showed that sparse recombination in 3.74% (+/−1.06 SEM) of the neurons. The vomeronasal neurons positive for tracing were variously distributed in apical and basal regions. Sox2 and tdTomato immunostaining revealed, as observed in the OE, that Islet1 tracing was detectable in sustentacular cells (Fig 7G). Since constitutive Isl1 lineage showed cells positive in olfactory and vomeronasal epithelia, we further investigated if Isl1 recombination also occurred in neurons of the septal organ and Gruenberg ganglion (GG). By performing immunostaining anti OMP in postnatal animals, we found that both neuronal populations were negative for lineage tracing. Immunostaining with anti-CART and -PDE2A antibodies also revealed an absence of Isl1 expression in Necklace glomeruli cells (NGCs) (data not shown).

**Fig. 7.**
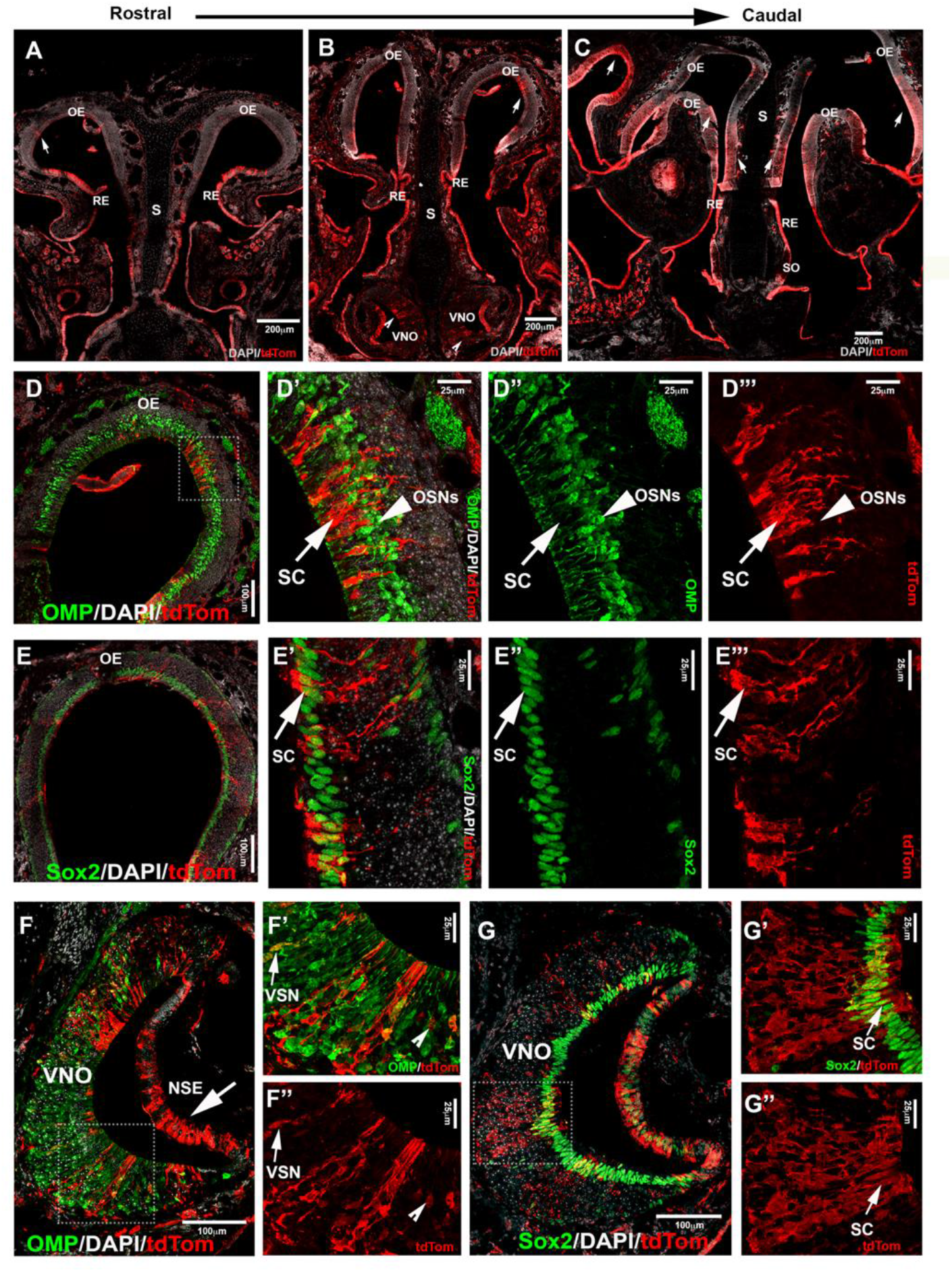
Lineage tracing of Isl1 positive cells postnatally highlights neuronal and non-neuronal cells in OE and VNE. **A,B,C**-Immunofluorescence against tdTom showing Isl1 tracing in olfactory epithelium (OE) from rostral toward the caudal ethmoid turbinates. Arrows (A,B,C) show tracing in OE. (B notched arrowheads) show tracing in VNO. **D-D’’’)** Double immunofluorescence against OMP (green) and tdTom (red) show no colocalization in neurons of the OE. **E-E’’’)** Double immunofluorescence against Sox2 (green) and tdTom (red) show Ils1Cre tracing (arrow marks) in the sustentacular cells of the OE. Sox2 is present in both globose basal cells that are present in the base of the OE and sustentacular cells in the upper layers of the OE. **F)** Double immunofluorescence against OMP (green) and tdTom (red) show Isl1 tracing in both non-sensory epithelium (NSE-arrow mark) and sensory epithelium of the VNO. **F’-F’’)** Arrow marks show colocalized mature vomeronasal neurons (VSN) and notched arrow heads show traced neurons that are not labelled with OMP. **G-G’’)** Double immunofluorescence against Sox2 (green) and tdTom (red) show Isl1 tracing in Sustentacular cells in the VNO.

### Conditional Isl1 loss of function in GnRH-1ns

Isl1 is important for differentiation, cell migration, survival and axonal targeting of neurons of the peripheral nervous system [53; 66; 67]. Our data revealed strong Isl1/2 expression in developing migrating GnRH-1ns (Fig.3, 3,4,6). To test if Isl1 plays a role in GnRH-1ns migration or controls GnRH expression, we generated GnRH-1Cre/Isl^flox/flox^ conditional mutants. Embryos were analyzed at E15.5, which is the stage at which the majority of the GnRH-1ns have already migrated in the brain [68]. Double immunostaining against Isl1/2 and GnRH confirmed detectable Isl1 ablation in 38% (SE +/−5.28%) of GnRH-1 neurons (Fig. 9 A-B’). While Isl1 in controls was detectable in virtually all migratory GnRH-1ns. In GnRH-1Cre/Isl^flox/flox^ conditional mutants, GnRHns negative for Isl1 immunoreactivity were found in the nasal area, forebrain junction and the brain. These data suggest that Isl1 expression is not required for GnRH-1 peptide expression nor GnRH-1 neuronal migration. Quantification of GnRH-1ns distribution in nasal area, forebrain junction and brain indicated an overall distribution of GnRH-1 neurons comparable with the one of controls (Fig 9. E).

## Discussion

Cellular derivatives of the olfactory placodes play central roles in respiration, chemosensory detection, sexual development, fertility and inter and intra species social behaviors. Understanding the basic mechanisms that define cellular diversity and underly normal and pathological development of the olfactory placode derivatives is of high clinical importance. Isl1 expression can occur in various neurogenic placodes [69; 70]; however, a comprehensive analysis of Isl1 expression remains unknown. Isl1, which is important for differentiation, cell migration, survival and axonal targeting of neurons in the peripheral nervous system [53; 66; 67], has a very high upregulation during GnRH-1 neuronal differentiation [52]. So, Isl1 remained a potential suitable genetic marker to label derivatives of the olfactory placode [49]. Here, we sought to trace Isl1 expression and lineage in the developing nasal area using in situ-hybridization, immunohistochemistry (Fig.3), temporally controlled Is1CreRT recombination (Fig. 4) and constitutive Isl1Cre genetic lineage tracing (Fig 5-8).

**Fig. 8.**
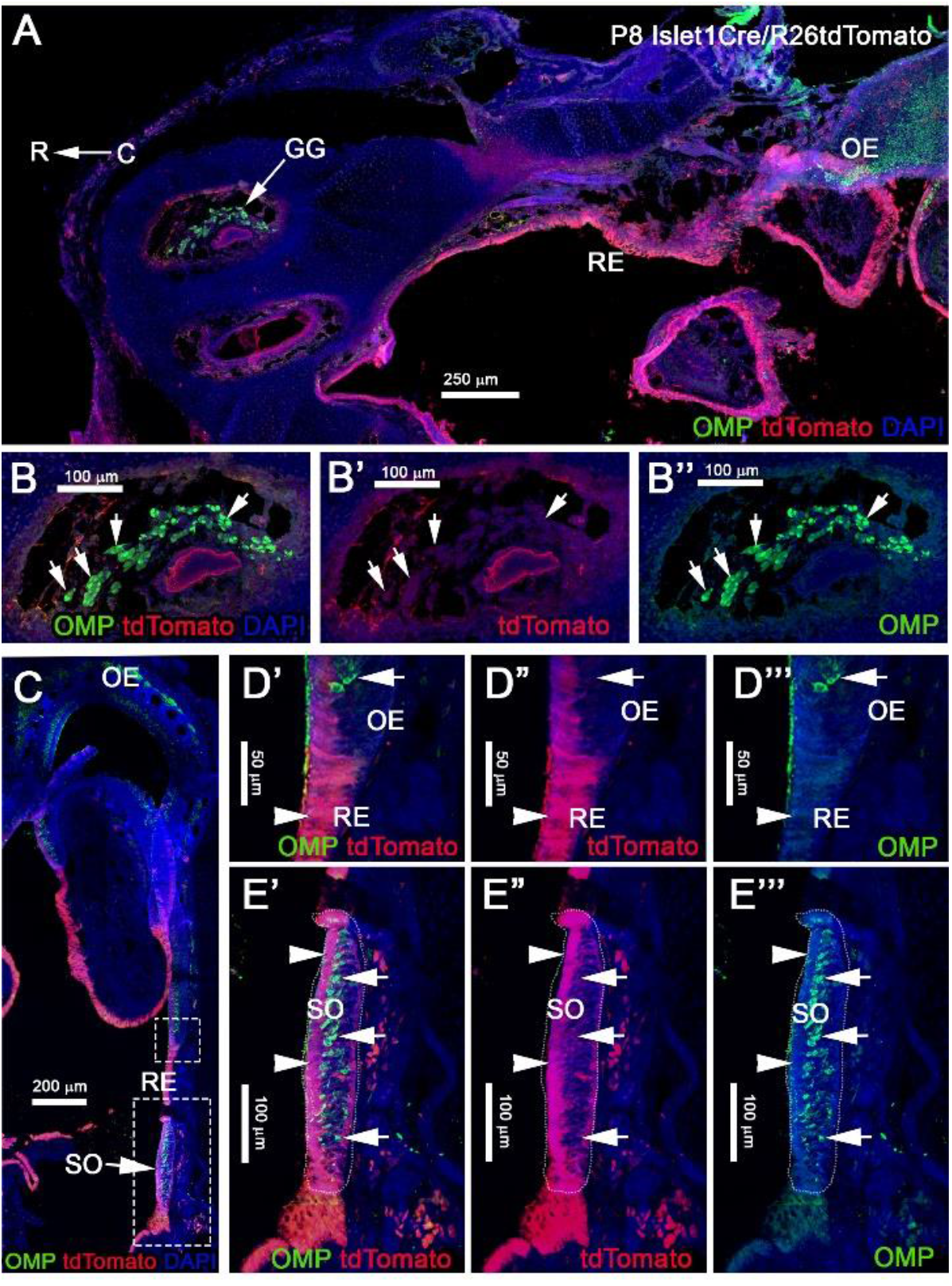
Post natal Isl1 constitutive lineage tracing is not expressed in other nasal sensory neurons. A) Isl1Cre/R26tdTomato P8 parasagittal section of the nose immunostained against OMP and tdTomato show that Isl1 recombination is limited to the respiratory epithelium even in post-natal stages. Neuronal sensory populations, such as the Gruenberg Ganglion GG B-B’’), is negative for recombination. C) Coronal view of Isl1Cre/R26tdTomato at P8, stained against OMP and tdTomato shows Isl1 tracing restricted to the non-sensory cells and cells of the respiratory epithelium. D-D’’) Magnification from C) of the border between olfactory and respiratory epithelium shows Isl1Cre tracing restricted to the respiratory epithelium (arrowhead), while OMP+ neurons are negative (arrow). E-E’’) Septal Organ (SO) from C) indicates Isl1 recombination is restricted to the non-sensory (arrowheads) cells, while OMP+ neurons are negative (arrows).

### GnRH-1 lineage tracing in the developing nasal area

Using in situ hybridization and Immunohistochemistry, we detected Isl1 expression at E11.5 in the putative GnRH neurogenic area, ventral to the developing VNO (Fig. 3A-D), in newly formed GnRH-1 neurons and the developing olfactory epithelium (Fig3D-F’’’). At E11.5, Isl1 immunoreactivity occurred in 89%(+/−3.33%) of GnRH cells, in line with published reports [49]. However, histochemistry at E13.5 and E15.5 showed strong Isl1/2 immunoreactivity in virtually all GnRH-1 neurons, suggesting that Isl1 expression is not limited to specific subpopulations of GnRH cells [49]. These data suggest a dynamic expression of Isl1 with more GnRH-1 cells positive across development.

At E11.5 we also found the existence of proliferative cells positive for Isl1 in the developing olfactory pit. Tamoxifen controlled Isl1Cre^ERT^ tracing at E11.5 showed Isl1 lineage in putative GnRH progenitors/newly formed GnRH-1 neurons, cells of the respiratory epithelium, few proliferative progenitors in the developing olfactory pit, neurons at the border between respiratory epithelium and developing olfactory epithelium and sparse neuronal (HuC/D+) cells in the OE and VNO. Notably in this experiment we analyzed Isl1Cre^ERT^/R26R recombination 4 days after Tam treatment. This means that, when we observed the histological samples, we identified either cells that expressed Isl1 at the moment of Tam injection or cells that derived from progenitors expressing Isl1 at time of Tam injection.

In line with what observed after temporally Tam controlled recombination at E11.5, constitutively active Isl1Cre recombination showed that, during embryonic development (Fig 5,6), Isl1 tracing highlights neurons in OE proximal to the border with RE, neurons in the vomeronasal organ, cells of the RE, GnRHns, migratory neurons negative for GnRH, and sustentacular cells in olfactory and vomeronasal organ.

These data indicate that Isl1 expression/lineage is limited to specific subsets of olfactory placodal derivatives. In contrast to prior observations during embryonic development, Isl1Cre recombination analysis of postnatal Isl1Cre/R26R mice showed only sparse neurons positive for Isl1 tracing in the most caudal regions of the olfactory sensory epithelium, while cells positive for Isl1 tracing were still found in the VNO, respiratory epithelium and in sustentacular cells of both OE and VNO (Fig.7). No Isl1 expression or lineage was in GC neurons and Grueneberg ganglion (Fig.8). These data suggest that Isl1 positive neurons in the OE at the border with the RE during embryonic development (Fig.4,5) either migrated out of the epithelium as a part of the migratory mass or died, even though we could still detect Isl1 expression/lineage in some vomeronasal organ neurons (Fig. 7 F-F’’). So, we further defined the fate of the Isl1 neurons detectable in the OE during embryonic development. During embryonic development we could also detect axonal projections positive for Isl1 traversing through the nasal area (Fig 4,5). Some axonal projections likely belonged to the nasopalatine projections of the trigeminal nerve (Fig 3, 4,5). Although both Isl1^CreERT^ and Isl1Cre induced recombination in Peripherin+ fibers projected to the ventral OB, the majority of OMP+ fibers projecting to the MOB and AOB appeared negative for Isl1 tracing (Fig 4,5). We did not pursue a detailed analysis of olfactory projections in postnatal animals, due to technical challenges in tracing Isl1 recombination in mitral cells of the MOB.

### Isl1 in the GnRH-1 system

During embryonic development GnRH-1ns migrate from the developing olfactory pit into the hypothalamus. Once in the preoptic area, GnRH neurons (GnRH-1ns) control the hypothalamic– pituitary–gonadal hormonal axis and reproductive development. Defective GnRH-1 release is the cause of Hypogonadotropic Hypogonadism, a condition that negatively impacts normal body development, social interactions, the ability to procreate, and physical performances [71; 72; 73; 74]. Hypogonadotropic Hypogonadism associated with congenital anosmia (impairment of the sense of smell) is clinically defined as Kallmann Syndrome [75; 76; 77; 78; 79; 80; 81]. Though postnatal development of the olfactory system has been extensively studied we still ignore many aspects regarding early development of the olfactory pit derivatives.

Previous studies in chick and mice indicate that soon after the olfactory placodes form they give rise to migratory olfactory pioneer neurons, and to several other neuronal populations including the terminal nerve cells and the GnRH-1ns. Placodal derived migratory neurons and the neural crest derived Olfactory ensheathing cells form the migratory mass [45; 82].

We previously shown that the fibers of the terminal nerve, upon which the GnRH-1ns migrate, are distinct from the olfactory and vomeronasal sensory neurons [28; 36; 83]. What progenitors give rise to the GnRH neurons, and what transcription factors control their onset are still open questions. Moreover, if what we define as GnRH neurons is one homogenous neuronal population is also a matter of debate.

Lineage tracing at different developmental stages suggest strong and consistent Isl1 expression in GnRH-1ns and cells of the migratory mass (Fig.3,4,6). While the terminal nerve, that projects from the vomeronasal area to the basal forebrain appeared to have heterogenous Isl1 expression/ genetic lineage tracing. (Fig. 6) Isl1 has been reported to be important in neuronal differentiation, cell migration, survival and axonal targeting of various neuronal types [53; 66; 67]. The consistent Isl1/2 expression in developing and migratory GnRH-1ns (Fig. 3,4,6), which is conserved from fish to humans prompted us to test its developmental role.

We first made an attempt to conditionally KO Isl1, which as a whole KO is embryonic lethal, by mating Isl1^CreERT^ mice (Fig. 4), which are Isl1^null^ heterozygous, with Isl1^flox^. After tamoxifen injection at E11.5 we analyzed Isl1^CreERT/flox^ mice and controls at E15.5 (as in Fig. 4). After Tamoxifen treatment Isl1^CreERT^ extensively recombines in the developing GnRH-1 neurons (Fig.4). Surprisingly Tam+ Isl1^CreERT^/Isl1^flox^ embryos did not show any obvious phenotype related to GnRH-1ns (supplementary data). However, for this model, we could not verify the efficiency of recombination because antibodies unable to detect the floxed Isl1 truncated protein were still recognizing the Isl1^CreERT^ protein product (see material and methods). For this reason, we report these as supplementary observations. In order to further test the role of Isl1 in GnRH-1 neuronal development we generated GnRH-1Cre/Isl1^flox/flox^ conditional mutants. Here, we could document complete loss of Is1 immunoreactvity in around ~40% of GnRH-1 neurons (Fig.9). Notably most of the anti Isl1 Abs have some level of cross reactivity with Isl2, therefore we performed in-situ hybridization against exon 4 of Isl1 paired with anti GnRH-1 immunofluorescent staining (not shown). These experiments suggest similar penetrance of recombination as indicated by IF. Notably, also in GnRH-1Cre/Isl1^flox/flox^ mutants we did not observe obvious defects in GnRH-1 neuronal migration nor in GnRH-1 expression (Fig. 9). In fact, GnRH-1 neurons negative for Isl1 were found in both nasal area and brain. Total number and overall distribution of GnRH-1 was comparable to controls. These data suggest that Isl1 alone is not necessary for GnRH-1 expression or neuronal migration. If Isl2 is also expressed in the GnRH-1ns and play a compensatory effect is worth further analysis.

**Fig 9.**
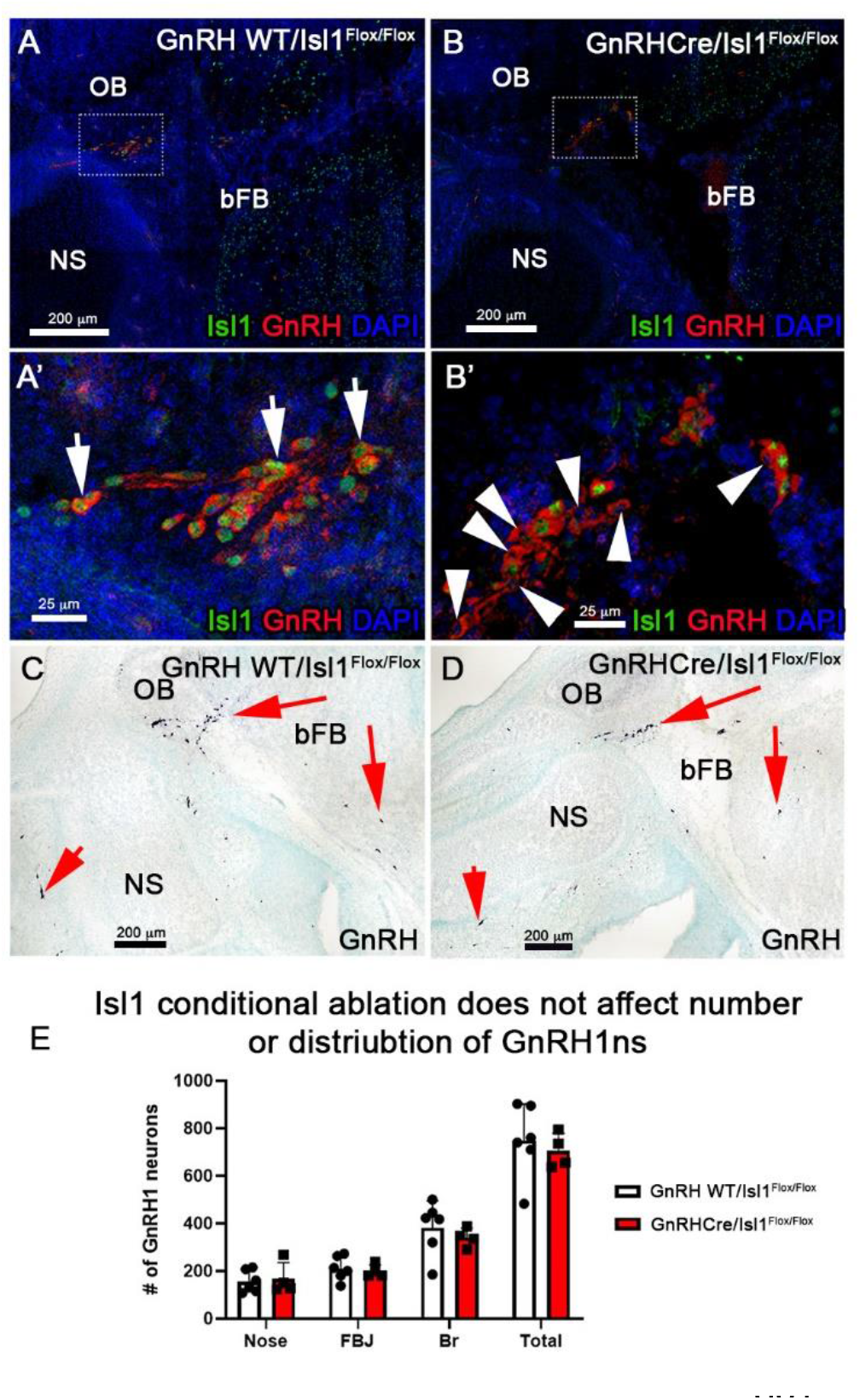
Conditional Loss of Isl1 in GnRHns has no effect. A-B’) Isl1/2 and GnRH double IF on E15.5 WT (A-A’), where all GnRHns express Isl1/2 (arrows) and cKO (B-B’) sections show that some GnRH-1ns lose the expression of Isl1 (arrowheads). C-D) Immunohistochemistry of GnRH-1 on WT and cKO sections show comparable distribution of cells positive for GnRH-1. E) Quantification of cKO show no changes in the number or distribution of GnRH-1ns. NS: Nasal Septum, OB: Olfactory Bulb, bFB: basal Forebrain

In conclusion, our data indicate very selective expression of Isl1 in neurons forming in the OP including the GnRH-1 neurons, various migratory neuronal populations and some cells in the VNO, terminal nerve and sustentacular cells on the more medial and lateral portions of the OE. Isl-1 immunoreactivity and genetic tracing suggest that Isl-1 expression, though dynamic, is common feature for virtually all GnRH-1 neurons [49] and several other neurons of the migratory mass. If all these early migratory neurons derive from similar pre placodal subregions should be further investigated. Performing GnRH-1ns specific Isl1 ablation in GnRHns we did not detect obvious cell specification, differentiation, survival or migratory defects [49] at embryonic development stages suggesting a dispensable role for Isl1 in GnRH neurons.

Our results leave open the question of what is the physiological role of this transcription factor in murine GnRH-1 neurons development and function. Further studies focused on Isl1 role in postnatal developmental and fertility can give definitive insights.

## Ethics Statement

All mouse studies were approved by the University at Albany Institutional Animal Care and Use Committee (IACUC).

## Author Contributions

EZT, set the mouse mating, collected samples prepared histological samples, performed immunostaining, imaging, and analyzed data at embryonic developmental stages. RRK prepared histological samples, performed immunostaining and analyses of postnatal animals. EZT and PEF designed the experiments wrote the article and prepared figure panels.

## Funding

Research reported in this publication was supported by the Eunice Kennedy Shriver National Institute of Child Health and Human Development of the National Institutes of Health under the Awards R15-HD096411 (PEF), and R01-HD097331/HD/NICHD (PEF) and by the National Institute of Deafness and other Communication Disorders of the National Institutes of Health under the Award R01-DC017149 (PEF).

**Supp. Fig. 1.**
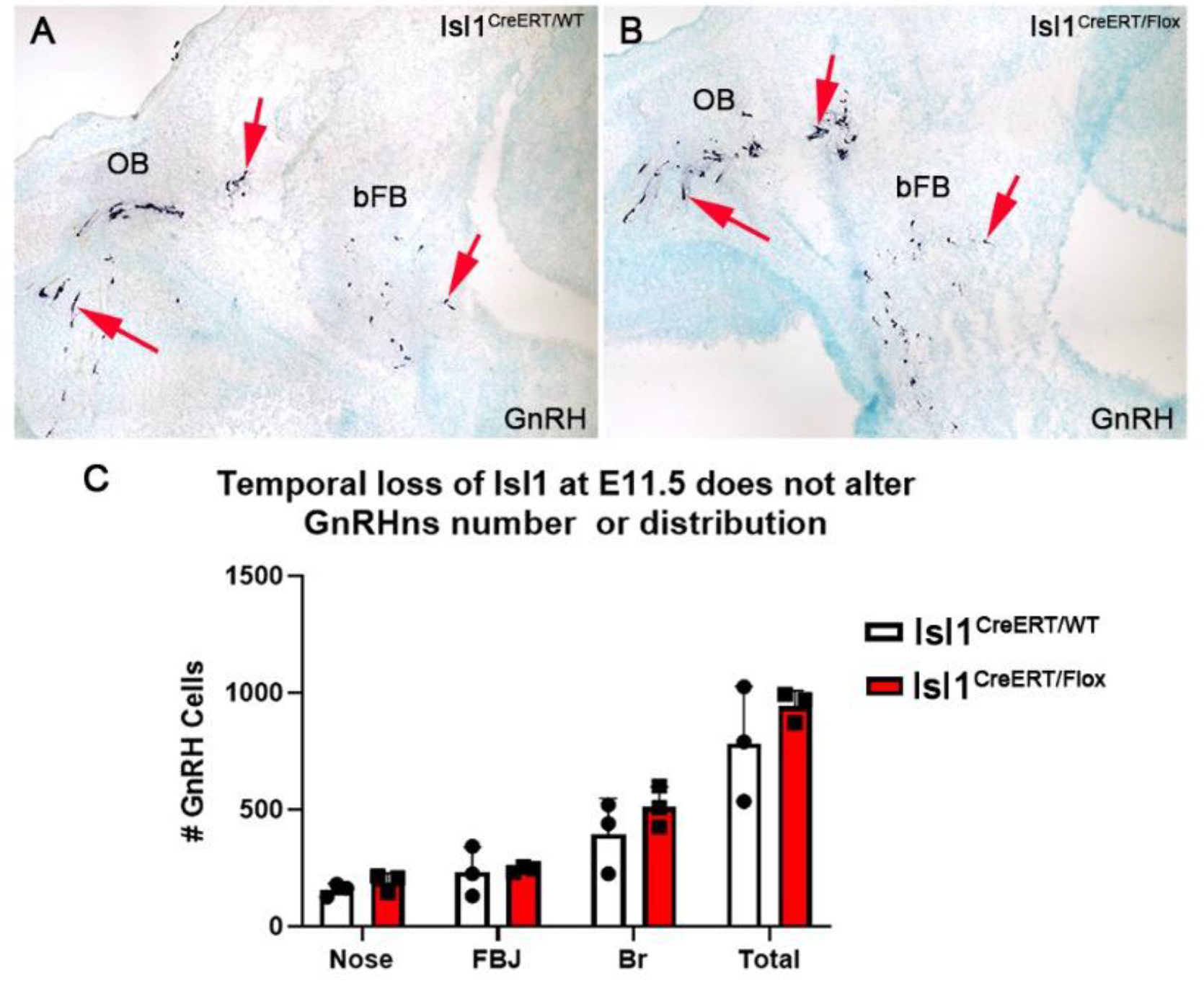
Conditional Loss of Isl1 at E11.5 does not affect GnRH1ns. A-B) GnRH IHC on WT(A) and cKO (B) sections show that between each genotype the distribution of GnRHns looks similar. C) Quantification of the number of GnRHns in whole embryo heads show no significant regional changes or in total number

